# Geographically Biased Composition of NetMHCpan Training Datasets and Evaluation of MHC-Peptide Binding Prediction Accuracy on Novel Alleles

**DOI:** 10.1101/2023.09.03.556092

**Authors:** Thomas Karl Atkins, Arnav Solanki, George Vasmatzis, James Cornette, Marc Riedel

## Abstract

Bias in neural network model training datasets has been observed to decrease prediction accuracy for groups underrepresented in training data. Thus, investigating the composition of training datasets used in machine learning models with health-care applications is vital to ensure equity. Two such machine learning models are NetMHCpan-4.1 and NetMHCIIpan-4.0, used to predict antigen binding scores to major histocompatibility complex class I and II molecules, respectively. As antigen presentation is a critical step in mounting the adaptive immune response, previous work has used these or similar predictions models in a broad array of applications, from explaining asymptomatic viral infection to cancer neoantigen prediction. However, these models have also been shown to be biased toward hydrophobic peptides, suggesting the network could also contain other sources of bias. Here, we report the composition of the networks’ training datasets are heavily biased toward European Caucasian individuals and against Asian and Pacific Islander individuals. We test the ability of NetMHCpan-4.1 and NetMHCpan-4.0 to distinguish true binders from randomly generated peptides on alleles not included in the training datasets. Unexpectedly, we fail to find evidence that the disparities in training data lead to a meaningful difference in prediction quality for alleles not present in the training data. We attempt to explain this result by mapping the HLA sequence space to determine the sequence diversity of the training dataset. Furthermore, we link the residues which have the greatest impact on NetMHCpan predictions to structural features for three alleles (HLA-A*34:01, HLA-C*04:03, HLA-DRB1*12:02).

## 1 Introduction

Antigen presentation by the major histocompatibility complex (MHC) class I and II proteins (referred to as HLA in humans) is one of the crucial steps to activating the adaptive immune response, and the genes which encode these proteins are some of the most polymorphic genes in humans [1]. As a result, the epitopes presented to T cells are determined partly by the binding affinity between the peptide fragment of the antigen and the host-specific MHC protein, which is determined by the amino acid sequences of both peptide and MHC. Because of the central role of this process in adaptive immunity, the ability to predict which peptides will bind to a given MHC allele has utility in diverse fields. For example, peptide-MHC binding predictions have been used to select peptides for a cancer neoantigen vaccine and to explain asymptotic SARS-CoV-2 infection in individuals with a specific HLA-B allele [2][3]. While molecular dynamics (MD) systems exists for modelling these complexes [4][5], the current consensus is that neural network prediction models are accurate enough at predicting binding affinity to be used in clinical settings [6]. Many such tools have been developed to predict peptide binding to both MHC class I and MHC class II [7][8][9][10]. Two of neural-network based predictors, NetMHCpan-4.1 and NetMHCIIpan-4.0 (here on out collectively referred to as NetMHCpan) are hosted on a popular web server and are fast to return predictions, making them popular choices for predicting peptide-MHC binding [11].

However, NetMHCpan does not rely on any structural information about the peptide or MHC molecule, and only takes an amino acid sequences for the peptide and MHC protein as input, which limits the model’s ability to generate mechanistic explanations for its binding predictions Additionally, the tool is closed-source, exacerbating its “black box” nature and prompting investigations into potential hidden biases. A previous study has shown NetMHCpan-4.1 has a previously unreported bias toward predicting hydrophobic peptides as strong binders, suggesting the predictions of these models need to be examined closely [12].

Many times when medical and biological neural network based prediction systems have been evaluated, researchers have uncovered numerous examples of racial bias in machine learning algorithms [13][14][15]. Furthermore, datasets from prior genomic studies often fail to capture the genetic diversity of the human population, often focusing on individuals of European descent, [16][17][18]. As these two significant effects intersect to produce models that overfit to overrepresented populations, it is vital that neural-network models be carefully investigated to determine the extent to which there is bias in the training dataset, and if it exists, the extent to which this bias affects the model predictions.

To determine the impact of training dataset bias on NetMHCpan’s predictions, we examined the geographic distribution of NetMHCpan’s training dataset and determined which populations are likely to have alleles not represented in NetMHCpan’s training dataset. We then measured the performance of NetMHCpan on alleles not present in its training dataset, and compared the performance to binding predictions for alleles present in its training dataset. To better understand these predictions, we created a map of HLA sequence space to determine the diversity of the dataset at the sequence level. Finally, for each of three MHC molecules not in NetMHCpan’s training dataset, we determined the residues of that molecule that have disproportionate impact on NetMHCpan’s predictions. This paper presents a geographic imbalance in the HLA types present in NetMHCpan’s training data, yet fails to find a significant drop in the accuracy of the network’s peptide binding predictions for alleles not present in the training data compared to the accuracy of the models’ prediction on alleles present in the training dataset. Furthermore, the results suggest two possible explanations for this finding. First, while the model may be lacking in geographic diversity, the alleles represented in the training dataset cover a large range of HLA sequences. Second, the model gives attention to residues structurally involved in peptide-MHC complexes for novel alleles.

## 2 Materials and Methods

### 2.1 MHC Allele Population Demographics

Data on HLA allele population frequencies were downloaded from the National Marrow Donor Program (NMDP) [19]. The dataset contains HLA-A/B/C/DRB1 population frequencies from *n* =6.59 million subjects from 21 self-reported racial groups, which are combined into six larger ethnicity categories, given in Supplementary Table S1. Because NetMHCpan uses a motif deconvolution algorithm for training, there exist data points in the eluted ligand dataset where a peptide corresponds to multiple MHC alleles [11]. In this case, we conservatively counted an allele as present in the training dataset if there is at least one positive example of a peptide binding to the associated cell line.

### 2.2. Evaluating NetMHCpan Performance

#### 2.2.1 Evaluation Datasets

In order to evaluate the performance of NetMHCpan, we used a dataset from Sarkizova et. al. [20]. The dataset consists of eluted ligand (EL) data for 31 HLA-A alleles, 40 HLA-B alleles, and 21 HLA-C alleles, with a median of 1,860 peptides per allele, generated by cell lines engineered to express only one HLA type. We excluded HLA-B alleles, as all forty of the HLA-B alleles had some presence in the NetMHCpan training data. We filtered the remaining peptides to only include 9-mers, and removed any 9-mers included in NetMHCpan’s training data from the evaluation set. Of these alleles, 7 (A*24:07, A*34:01, A*34:02, A*36:01, C*03:02, C*04:03, and C*14:03) have no representation in NetMHCpan’s training data (binding affinity or eluted ligand).

As no similar dataset exists for MHC class II, we created an evaluation set by downloading peptides from IEDB [21]. For each allele, the filters used were “Include Positive Assays”, “No T cell assays”, “No B cell assays”, and “MHC Restriction Type: [allele] protein complex.” To choose DRB1 alleles of interest, we selected alleles for which NetMHCIIpan-4.0 had eluted ligand data from a cell line engineered to express only one HLA-DRB1 allele. To obtain data for HLA-DRB1*12:02, we use a eluted ligand dataset from cell line C1R expressing HLA-DR12/DQ7/DP4 [22]. Because the cell line expressed both HLA-DRB1*12:02 and HLA-DRB3*02:02:01, Gibbs Cluster was used to separate the two groups [23] (Supplementary Figure S1). The group belonging to DRB1*12:02 was identified by the absence of F at P1, the absence of N at P4, and the presence of Y/F at P9.

To provide negative controls for both MHC class I and II, the real peptides were combined with randomly generated peptides so that the ground truth peptides made up 1% of the final evaluation set. For the MHC class II dataset, the length distribution of the randomly generated peptides was fixed to be equal to the length distribution of the ground truth peptides. Peptides were generated by choosing each amino acid at random with frequencies corresponding to amino acid frequencies in the human proteome.

#### 2.2.2 Log Rank Predictions, Motif Entropy Correction, and AUC

As a result of the above preprocessing steps, we obtained a dataset for 31 HLA-A alleles, 21 HLA-C alleles, and 11 HLA-DRB1 alleles, each dataset being made up of 1% peptides experimentally verified to bind to the HLA allele in eluted ligand assays, and 99% randomly generated peptides (to serve as a control). Random peptides are generated by randomly sampling amino acids using all organism amino acid frequencies [24]. Testing the methods with randomly generated peptides sampled directly from the human proteome did not significantly change the results (Supplementary Figure S2). For each allele, we used NetMHCpan-4.1 or NetMHCIIpan-4.0 to generate an eluted ligand score for each peptide in the training dataset, and ranked all peptides by their EL scores. We then measured performance based on the distribution of log ranks for the experimentally verified peptides. For example, if the model is a perfect predictor, all real peptides will have a log_10_ rank below −2, and if the model is a random predictor, 90% of real peptides will have a log_10_ rank between 0 and −1.

To correct for any discrepancies in difficulty predicting ligands based on selectivity of the MHC binding motif, we calculated the Shannon entropy of the binding motif for each allele as *−* _*a*_ *p*_*a*_ log_2_(*p*_*a*_), where *p*_*a*_ is the frequency of amino acid *a* in the allele-specific experimentally verified binding peptides. We then performed a linear regression for the log-rank against the entropy, shown in Supplementary Figures S3 and S4. For both MHC class I and class II, we found alleles with lower entropy (more predictable) motifs were associated with better predictions, as expected. Therefore, we created a correction factor for each allele measuring the expected difference in predictions compared to the mean, and subtracted that from the distributions to be able to compare alleles with different binding motif entropies. Additionally, because MHC class II proteins bind a core motif that can contain additional amino acids on the ends that do not affect the binding prediction, we encountered cases in the MHC class II dataset where multiple versions of a peptide contained the same core seqeunce, with minor discrepancies at the start and end of the peptide. Therefore, in this case, we chose to weight the MHC class II peptides based on NetMHCIIpan-4.0’s reported binding core, such that each core was weighted equally.

To determine a 95% confidence interval for the difference between the median of the ranks of the alleles with and without training data, a bootstrap procedure was used. Data were sampled with replacement for a number of times equal to the size of the data, and the difference between the medians of the bootstrap samples was calculated. This was repeated 10^6^ times, and the 0.025 and 0.975 quantiles were reported as the 95% confidence interval.

Finally, we calculate AUC as the area under the ROC (TPR-FPR) curve. The true positive rate (TPR) is defined by *TPR* = *TP/*(*TP* + *FN*), and the false positive rate (FPR) is defined by *FPR* = *FP/*(*FP* + *TN*). True positives are defined as experimentally verified peptides with a score greater than a given cutoff, and false positives as randomly generated peptides with a score greater than a given cutoff. True negatives are defined as as randomly generated peptides with a score less than a give cutoff, and false negatives as experimentally verified peptides with a score less than a given cutoff.

### 2.3 MDS of HLA Alleles

Using the NMDP frequency database, HLA-A, B, C, and DRB1 alleles with a frequency greater than 0.01% in any population were selected (*n* = 135 HLA-A, *n* = 258 HLA-B, *n* = 66 HLA-C, *n* = 118 HLA-DRB1). The IPD-IMGT/HLA alignment tool was used to create an alignment of the selected HLA full protein sequences [25]. In cases where large gaps occurred at the beginning or end of the alignment, gaps were filled with the most common amino acid occurring at that residue. Similarity between sequences was measured by summing the values of the PAM100 matrix for each pair of amino acids in the two sequences [26]. Distance was then measured as the difference between the maximum similarity and the computed similarity, normalized so that the maximum distance was reported. Scikit-learn’s MDS algorithm with default parameters was used to compute the MDS [27][28].

### 2.4 NetMHCpan Residue Substitution Sensitivity

Here, we describe a technique similar to the occlusion sensitivity technique common in the field of computer vision. We chose the alleles HLA-A*34:01, HLA-C*04:04. and HLA-DRB1*12:02 for the following experiments, as NetMHCpan performed the poorest on these three alleles. For each allele, we used NetMHCpan to predict the eluted ligand score for all the experimentally verified peptides, using an unmodified version of the MHC sequence. Next, for residues 1-205 (29-125 for DRB1*12:02), we asked NetMHCpan to predict the eluted ligand score for all experimentally verified peptides, using a version of the MHC sequence where for each residue, each of the other 19 amino acids was substituted. From this, we took the 5 amino acids for which NetMHCpan predicted the lowest scores, and calculated the mean difference between EL scores for the mutated and unmutated predictions, as to investigate the effect of replacing residues with dissimilar amino acids. Repeating this for every residue, we then obtained a metric for the relative importance of the residue to NetMHCpan’s predictions. HLA tertiary structures were generated using PANDORA and visualized using PyMOL [4], [29].

### 2.5 Software Versions

The following software versions were used: NetMHCpan (4.1), NetMHCIIpan (4.0), PANDORA (2.0), GibbsCluster (2.0), PyMol (2.6.0a0), sklearn (1.3.0). For any software that had options for a web-based and local version, a local version was always used.

## 3. Results

### 3.1 Common European Caucasian HLA Types are Overrepresented in NetMHCpan Training Data

As neural network prediction biases are often enforced by disparities in the amount of model training data, we first investigate NetMHCpan’s training dataset to determine whether the data is representative of the global population. To do this, we used allele distribution data from the National Marrow Donor Project (NMDP) [19]. Codes for population groups can be found in Supplementary Table S1. For each population, we calculated the fraction of people who have at least one HLA-A/B/C/DRB1 allele for which there is no data in NetMHCpan’s training set.

There exists a substantial disparity between the most and least represented populations in NetMHCpan’s training dataset. European Caucasian individuals are most likely to see their genotypes represented in the training set, while Southeast Asian, Pacific Islander, South Asian, and East Asian individuals are least likely to have genotypes represented in the training set (Figure 1). Using the NMDP categories, only 0.4%/0.9%/0.6%/2.6% of European Caucasian individuals have an HLA-A/B/C/DRB1 allele not found in NetMHCpan’s training data, while 5.1%/27.7%/12.1%/33.6% of Vietnamese individuals and 30.1%/39.3%/10.8%/16.1% of Filipino individuals have an HLA-A/B/C/DRB1 allele not found in NetMHCpan’s training data.

**Figure 1.**
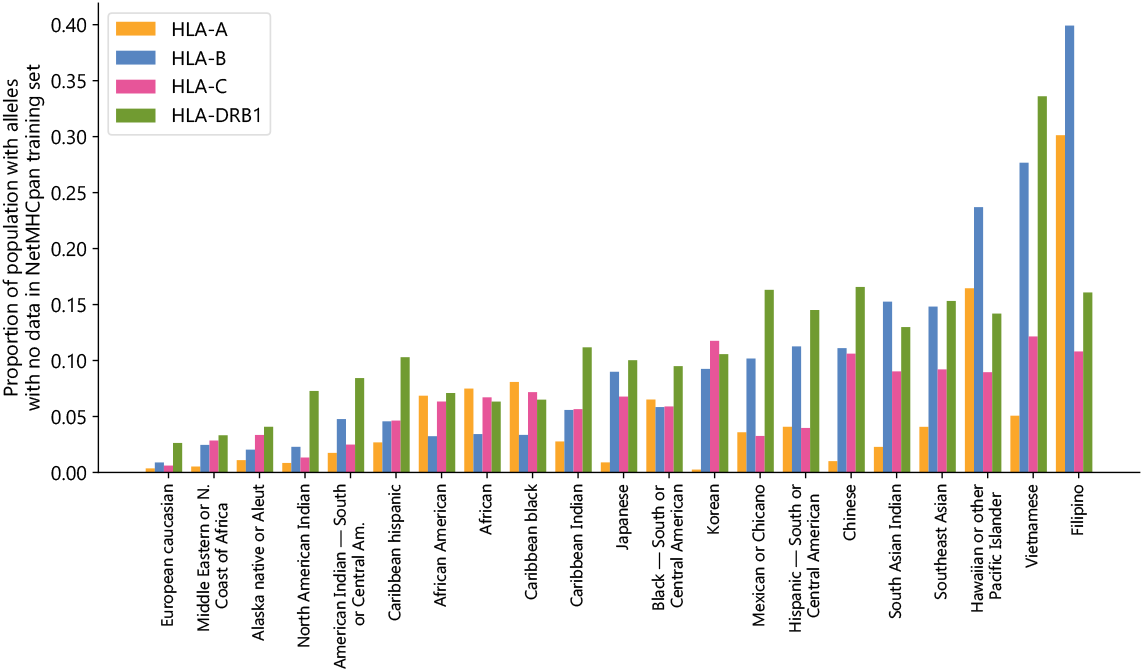
NetMHCpan training data fails to cover common HLA alleles: Proportion of populations (as defined by the National Marrow Donor Program) that have at least one HLA class A, B, C, or DRB1 allele with no data in the NetMHCpan-4.1 or NetMHCIIpan-4.0 training datasets.

These disparities are not likely to have arisen by chance alone, given the fractions of the populations for which no data exists are correlated between HLA groups (Supplementary Table S2). For all pairs of groups there exists a positive correlation, with the strongest correlation between HLA-A and HLA-B (0.750) and the weakest correlation between HLA-A and HLA-DRB1 (0.238). Because the disparities are found in all four HLA groups examined and are correlated with each other, this suggests a common systemic factor driving the extreme imbalance of the training dataset.

### 3.2 NetMHCpan-4.1 and NetMHCIIpan-4.0 Accurately Predict Peptide Binding to Novel Alleles

Because there exists such a vast disparity in the representation of populations in NetMHC-pan’s training data, we hypothesized NetMHCpan is overfitting to the training set, making the model unable to make accurate predictions for peptides binding to novel MHC proteins. Therefore, we investigated whether there is a decrease in prediction quality for HLA sequences not found in the training data. To do this, we performed an experiment in which NetMHCpan was tasked to predict eluted ligand binding scores for a dataset consisting of 1% peptides experimentally verified to bind to their corresponding MHC proteins and 99% randomly generated peptides, as has been commonly used in the literature [30]. We then measured the rank of the predictions for the experimentally verified peptides, which we use as our metric for the accuracy of the predictions (after a correction for motif binding entropy described in the Methods section), as well as the area under the ROC curve for each set of predictions (AUC).

We ran the MHC class I peptide experiment on a large HLA class I eluted ligand dataset [20]. In the dataset are *n* = 39617 peptides for 27 HLA-A and 18 HLA-C alleles with training data in NetMHCpan-4.1’s training set, and *n* = 8652 peptides for 4 HLA-A alleles and 3 HLA-C alleles without data in NetMHCpan-4.1’s training set. All together, these novel alleles represent up to 28.8% of HLA-A alleles, and up to 11.7% of HLA-C alleles for some populations (Supplementary Figure S5). Because there are no HLA-B alleles present in the dataset but absent from NetMHCpan-4.1’s training set, we omit HLA-B from this analysis.

NetMHCpan-4.1 accurately recalls experimentally validated peptides from a training dataset containing validated peptides and randomly generated peptides for these 7 alleles. For both HLA-A and HLA-C, the allele for which NetMHCpan-4.1 best recalled experimentally validated peptides was an allele for which NetMHCpan-4.1 had no data in its training set (A*24:07 and C*14:03) (Figure 2). Overall, the predictions of binding peptides for the alleles for which NetMHCpan-4.1 has no training data slightly outperform the predictions for alleles for which it does have data (Supplementary Figure S6), with a 95% bootstrap confidence interval for the difference in the medians of the two sets being (0.037, 0.063) (Supplementary Figure S7). On average, NetMHCpan-4.1 ranks experimentally verified peptides for alleles for which data does not exist 1.12 times higher than it ranks peptides which correspond to alleles in its training dataset. In summary, we fail to find evidence that the imbalance in the training dataset leads a decrease in the quality of NetMHCpan-4.1 predictions for novel alleles.

**Figure 2.**
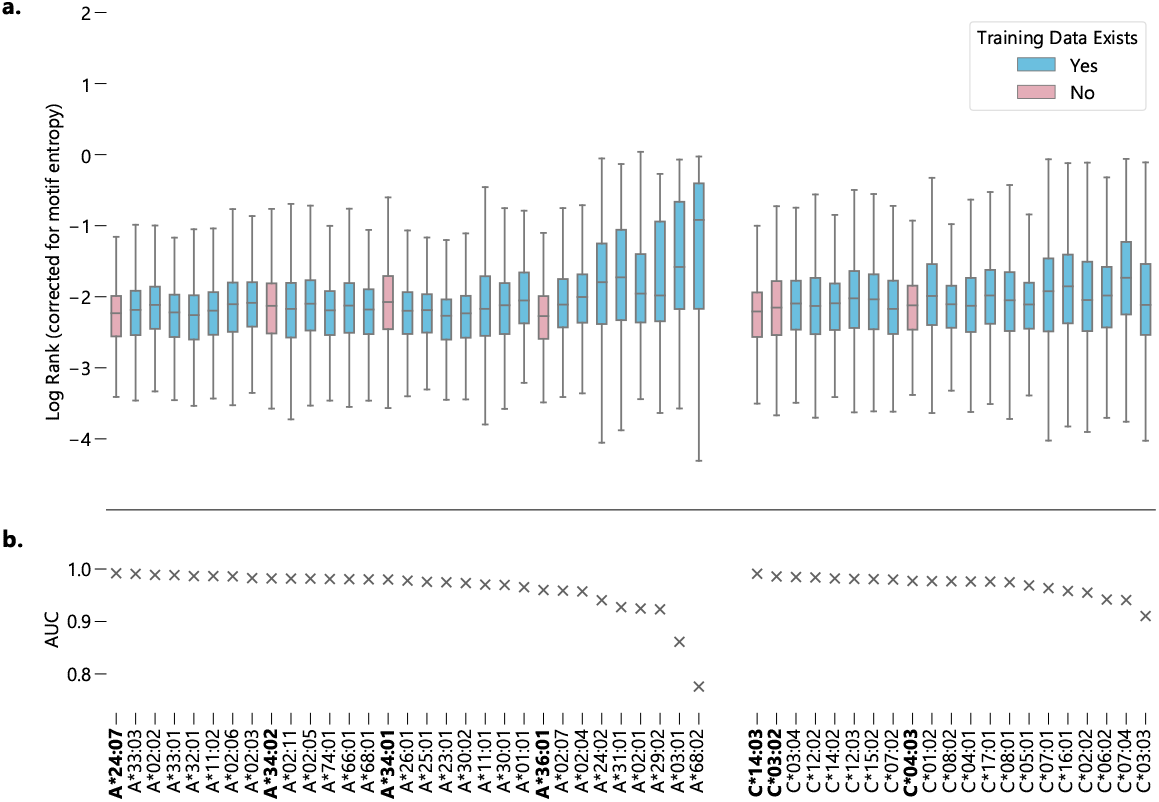
Evaluating NetMHCpan-4.1 performance on novel alleles: NetMHCpan-4.1 was tasked with separating peptides identified as true binders using LC-MS/MS (from Sarkizova et. al.) from randomly generated peptides for 52 HLA class I alleles. (A) Alleles with training data in NetMHCpan-4.1’s training dataset are shown in blue, alleles without are shown in pink. Performance is measured by the distribution of log ranks of the true peptides, corrected for entropy of the allele binding motif (lower is better). (B) Area under the ROC curve (AUC) for each allele.

**Figure 3.**
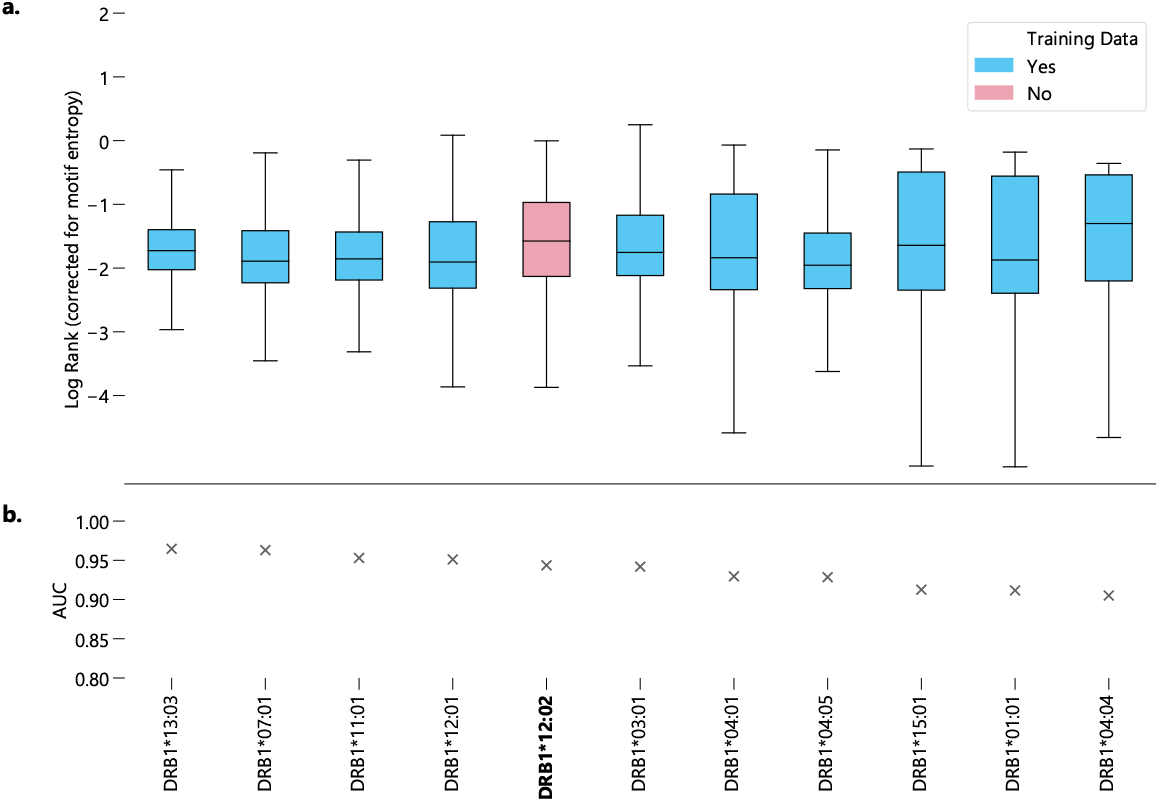
Evaluating NetMHCIIpan-4.0 performance on novel alleles: NetMHCIIpan-4.0 was tasked with separating peptides identified as true binders using LC-MS/MS (from IEDB) from randomly generated peptides for 10 HLA-DRB1 alleles with data in NetMHCIIpan-4.0’s training set, and one allele without training data. (A) Performance is measured by the distribution of log ranks of the true peptides, corrected for entropy of the allele binding motif (lower is better). (B) Area under the ROC curve (AUC) for each allele.

In the case of MHC class II predictions, we focus exclusively on DRB1 because HLA-DR is the only MHC class II protein to vary only in the beta chain, which simplifies the testing process, as we do not have to test combinations of alleles. While a comprehensive eluted ligand dataset exists for the MHC class I peptidome, no analogous dataset exists for HLA-DRB1. Therefore, we used IEDB to gather data for alleles which were present in NetMHCIIpan-4.0’s training data, and data from a recent C1R cell line eluted ligand study for peptides binding to DRB1*12:02, an allele not represented in NetMHCIIpan-4.0’s training set [21][22]. All together, we have *n* = 45286 peptides from 10 alleles with training data in NetMHCIIpan-4.0, and *n* = 32402 peptides from allele DRB1*12:02.

In contrast to NetMHCpan-4.1, the predictions generated by NetMHCIIpan-4.0 for peptides corresponding to alleles for which it has no data are slightly worse than average, when measured by median log-rank (Supplementary Figure S6). However, when measured by AUC, DRB1*!2:02 ranks around average, greater than 6 alleles and less than 4 alleles. A 95% bootstrap confidence interval for the difference in the medians between peptides corresponding to alleles with and without data in NetMHCIIpan-4.0’s training set is (−0.260, −0.232), with NetMHCIIpan-4.0 on average ranking experimentally validated peptides 1.8 times lower for the DRB1*12:02 allele (Supplementary Figure S7). However, while NetMHCIIpan-4.0 makes statistically significantly worse predictions for DRB1*12:02 than for the other alleles, the discrepancy between the median log rank of the best performing allele (DRB1*15:01) and the median log rank of DRB1*12:02 is less than half the interquartile range of the log-ranks of predictions for DRB1*12:02, suggesting the the difference in prediction quality is relatively minor compared to the variability in predictions for a given allele. Furthermore, there exists an allele with data in NetMHCIIpan-4.0’s training dataset, DRB1*04:04, for which NetMHCIIpan-4.0 is less accurate at distinguishing real peptides than for DRB1*12:02.

While problems of skewed datasets have affected quality of numerous other machine learning based predictions algorithms, we find no evidence this is true of NetMHCpan. By testing the ability of NetMHCpan to recall experimentally verified binding peptides to alleles for which the algorithm has no training data, we fail to conclude there exists a meaningful difference between alleles for which NetMHCpan has training data, and those for which it does not.

### 3.3 NetMHCpan Training Data Covers a Large Subset of HLA Allele Space

As a lack of diversity in training data often leads machine learning models to overfit to their training set, we seek to understand why this does not appear to be true for NetMHCpan. Therefore, we visualize the training dataset by measuring sequence similarity between HLA alleles with frequency greater than 0.01% in any population, and use these computed similarities to perform multidimensional scaling (MDS) in order to visualize the sequence space as a two-dimensional map [28].

For all four HLA types measured, alleles tend to organize into clusters, a majority which contain at least one allele with data in NetMHCpan’s training dataset (Figure 4a). This suggests that while NetMHCpan may be missing data for many alleles common in non-European populations, the alleles for which it has data are sufficiently similar to the missing alleles as to allow the model to make reasonable inference about the biochemical properties of alleles without data.

**Figure 4.**
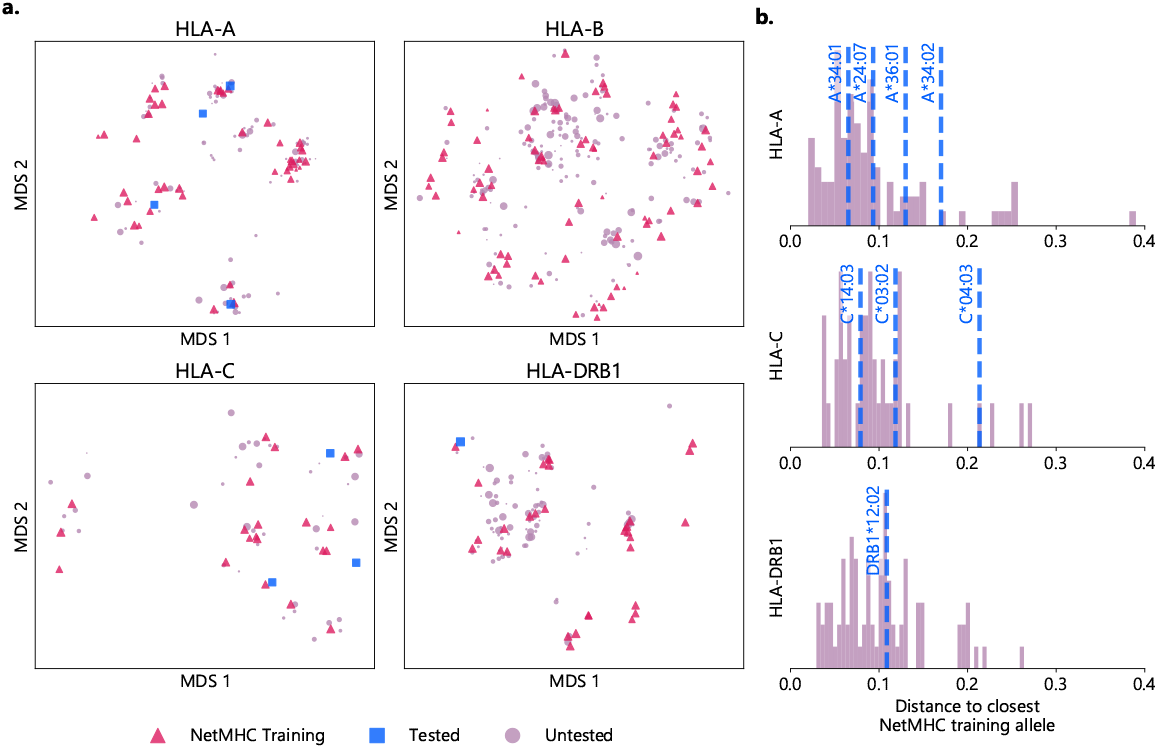
Visualizing the training space of NetMHCpan: (A) MDS plot of HLA alleles, with smaller distances corresponding to greater sequence similarity. Alleles included in NetMHCpan’s training data are marked with pink triangles, alleles tested in Figures 2 and 3 with no training data are marked with blue squares, and other alleles are marked with purple circles. Marker size corresponds to maximum frequency of the allele in any NMDP population (log scale). (B) Histogram of distance to closest allele to data in NetMHCpan’s training set for all alleles without training data. Alleles previously tested are shown with vertical dashed blue lines.

Furthermore, measuring pairwise distances between all alleles provides context for the performance of NetMHCpan on novel alleles reported above. While a sample size of *n* = 8 is not large enough to provide numerical estimates with any sort of power, the data qualitatively indicate a potential positive correlation between distance to the nearest allele and performance (Supplementary Figure S8). To measure the extent to which an allele is novel, we calculate the distance to the nearest allele in the training data for each allele not in NetMHCpan’s training data (4b). Of the eight alleles tested, seven are further from the nearest allele present in training data than a majority of the untested alleles, with the exception being C*14:03 (Supplementary Table S3). Therefore, while the choices of which alleles without training data to test were driven by data availability, we demonstrate the alleles tested are less similar to the training data than other HLA alleles. Thus, the accuracy of NetMHCpan’s predictions for these alleles is not driven by greater than average similarity of these alleles to alleles found in the training dataset.

### 3.4 NetMHCpan Correctly Identifies MHC Residues Involved in Peptide Binding

Finally, we aim to understand the extent to which NetMHCpan identifies residues structurally involved in peptide binding. As NetMHCpan allows for direct input of an MHC protein sequence, we perform an experiment in which we mutate each residue of a given HLA sequence individually, and measure how much NetMHCpan’s EL scores for experimentally verified peptides change compared to the unmodified sequence. We focus on three case studies, HLA-A*34:01, HLA-C*04:03, and HLA-DRB1*12:02, as these alleles constitute the worst-performing allele for each HLA type.

In each case, the MHC residues which have the greatest impact on NetMHCpan’s prediction are all residues that make physical contact with the peptide (Figure 5, Supplementary Tables S4-S6). This suggests that the accuracy of NetMHCpan’s predictions on novel alleles is partly driven by its ability to selectively pay attention to residues involved with the physical process of binding. Of special interest is the observation that many residues which affect the predictions for peptides binding to DRB1*12:02 are residues previously identified to determine the binding motif of DR12, namely, 13G, 57V, 70D, 71R, 74A, and 86V [22]. Therefore, we conclude NetMHCpan implicitly learns the MHC residues structurally involved in binding, and its ability to generalize these findings to novel alleles increases its prediction accuracy.

**Figure 5.**
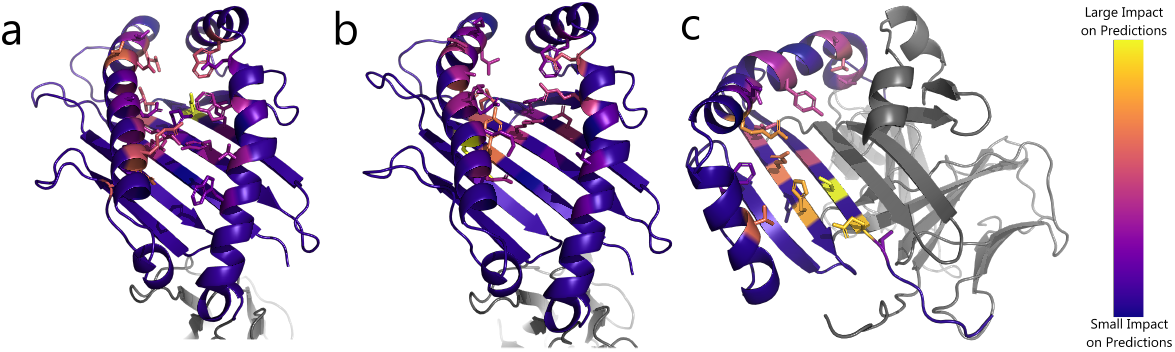
Impact of substituting residues on NetMHCpan predictions for HLA alleles of interest: Structure of (A) HLA-A*34:01 (B) HLA-C*04:03 and (C) HLA-DRB1*12:02. Residues are colored by impact of substitution on NetMHCpan predictions. Yellow resides indicate a large change to NetMHCpan predictions when replaced, purple resides indicate a small change. Sidechains are shown for residues of interest

## 4 Discussion

We report NetMHCpan fails to include a geographically diverse set of HLA alleles in its training data. We find individuals from underrepresented populations, predominantly from Asia, are twenty times more likely to carry HLA alleles not present in NetMHCpan’s training data. Furthermore, we observe correlation between population representation between all four alleles measured, suggesting that the dataset bias is a result of systemic underrepresentation of minority groups in the NetMHCpan training dataset.

Numerous previous examples of training dataset racial bias affecting machine learning model predictions led us to hypothesize NetMHCpan would make less accurate predictions on alleles which were not present in its training dataset [13][14][15]. Furthermore, previous work showed NetMHCpan is subject to systemic biases regarding hydrophobicity, suggesting that other biases may be lurking [12]. Unexpectedly, we fail to find evidence that there is a substantial difference in the ability of NetMHCpan to discriminate experimentally verified binding peptides from randomly generated peptides. Instead, we observe a slight increase in the prediction ability for MHC class I alleles with no data present in the training set, and only a slight decrease for MHC class II alleles. While both effects are statistically significant, we allege neither is large enough to have a substantial effect on prediction quality.

To explain this unexpected result, we characterize the sequence space of common HLA alleles. While NetMHCpan’s training dataset fails to include many alleles common in underrepresented populations, we show that the alleles for which training data exist are well-distributed throughout sequence space. We thus hypothesize that MHC sequence diversity in the training dataset partially explains the failure to observe a drop in prediction quality.

Furthermore, we establish a connection between the residues that impact NetMHCpan’s predictions and the residues that physically contact the peptide for three HLA alleles not present in NetMHCpan’s training data.

The discrepancies in the diversity of HLA eluted ligand datasets that compelled this study also constitute a major limitation, as only eight novel HLA alleles were tested, with no novel HLA-B alleles. Furthermore, our study design was limited to only testing one allele at a time, and so we did not investigate complex effects that could be associated with linkage disequilibrium in MHC class II molecules formed by two interacting chains, including HLA-DQ and HLA-DP [31]. We only tested the ability of NetMHCpan to distinguish experimentally verified peptides from randomly generated peptides, and did not perform any experiments to characterize the model’s ability to predict binding affinity. Finally, NetMHCpan is closed source, and so we were unable to view the internal network structure, needing to rely on an occlusion sensitivity-like metric to determine how the network makes predictions.

We present evidence of a strong bias in NetMHCpan’s training dataset toward European Caucasian individuals. While we fail to find evidence this bias affects the accuracy of NetMHCpan’s predictions, the bias in the training dataset highlights the need for MHC eluted ligand datasets that contain data for alleles for underrepresented populations. Furthermore, given the outsized impact of NetMHCpan on the training data generated for other MHC binding prediction tools, future work must investigate the composition of training datasets and potential bias in other tools [32]. Finally, we recommend all tools that utilize a dataset involving HLA alleles as part of their pipeline clearly report the composition of any datasets they utilize for training, and perform additional testing in the presence of biased training data to ensure model predictions do not substantially decline for underrepresented groups.

## Author Contributions

TKA, AS, GV, JC, and MR led discussions on this research. TKA conducted the data analyses and wrote the manuscript. JC, GV, and MR reviewed the manuscript. All authors contributed to the article and approved the submitted version.

## Acknowledgments

We thank Mohammed Yaro for sharing his expertise in the field of MHC.

## Supplemental Data

Supplementary Material should be uploaded separately on submission, if there are Supplementary Figures, please include the caption in the same file as the figure. LaTeX Supplementary Material templates can be found in the Frontiers LaTeX folder.

## Data Availability Statement

The datasets generated for this study can be found in the following respository: https://github.com/ThomasKAtkins/netmhcpan-bias-data/tree/main.

