## Supplementary Tables and Figures for "Geographically Biased Composition of NetMHCpan Training Datasets and Evaluation of MHC-Peptide Binding Prediction Accuracy on Novel Alleles"

### 1 Supplementary Tables and Figures

#### 1.1 Tables

| Code | Description | Group |
|---|---|---|
| AAFA | African American | AFA |
| AFB | African | AFA |
| AINDI | South Asian Indian | API |
| AISC | American Indian — South or Central Am. | NAM |
| ALANAM | Alaska native or Aleut | NAM |
| AMIND | North American Indian | NAM |
| CARB | Caribbean black | AFA |
| CARHIS | Caribbean hispanic | HIS |
| CARIBI | Caribbean Indian | NAM |
| EURCAU | European caucasian | CAU |
| FILII | Filipino | API |
| HAWI | Hawaiian or other Pacific Islander | API |
| JAPI | Japanese | API |
| KORI | Korean | API |
| MENAFC | Middle Eastern or N. Coast of Africa | CAU |
| MSWHIS | Mexican or Chicano | HIS |
| NCHI | Chinese | API |
| SCAHIS | Hispanic — South or Central American | HIS |
| SCAMB | Black — South or Central American | AFA |
| SCSEAI | Southeast Asian | API |
| VIET | Vietnamese | API |

Table S1: Full descriptions for codes given by NMDP project

| | A | B | C | DRB1 |
|---|---|---|---|---|
| A | - | 0.750 | 0.401 | 0.238 |
| B | | - | 0.684 | 0.732 |
| C | | | - | 0.643 |
| DRB1 | | | | - |

Table S2: Correlations between absence of HLA alleles in population groups

| Allele | Rank | Closest Allele |
|---|---|---|
| A*24:07 | 0.648 | A*24:02 |
| A*34:01 | 0.571 | A*66:01 |
| A*34:02 | 0.901 | A*66:01 |
| A*36:01 | 0.813 | A*01:01 |
| C*03:02 | 0.733 | C*03:04 |
| C*04:03 | 0.911 | C*04:01 |
| C*14:03 | 0.356 | C*14:02 |
| DRB1*12:02 | 0.581 | DRB1*12:01 |

Table S3: Fraction of alleles that have a closer distance to an allele in NetMHCpan's training set than the given allele

| Pos | AA | Change |
|---|---|---|
| 33 | Y | 0.157 |
| 48 | A | 0.039 |
| 69 | M | 0.230 |
| 86 | R | 0.218 |
| 87 | N | 0.260 |
| 90 | K | 0.155 |
| 91 | V | 0.109 |
| 93 | A | 0.080 |
| 94 | Q | 0.107 |
| 97 | T | 0.064 |
| 98 | D | 0.153 |
| 100 | V | 0.262 |
| 101 | D | 0.194 |
| 104 | T | 0.107 |
| 105 | L | 0.219 |
| 119 | I | 0.215 |
| 121 | R | 0.063 |
| 123 | Y | 0.123 |
| 138 | Q | 0.113 |
| 140 | D | 0.420 |
| 167 | T | 0.025 |
| 171 | W | 0.210 |
| 176 | E | 0.083 |
| 180 | W | 0.137 |
| 182 | A | 0.034 |
| 187 | T | 0.123 |
| 191 | W | 0.072 |

Table S4: Average change in NetMHCpan-4.1 EL score for experimentally binding peptides under residue substitution for peptides binding to HLA-A*34:01

| Pos | AA | Change |
|---|---|---|
| 33 | Y | 0.182 |
| 48 | A | 0.032 |
| 86 | R | 0.017 |
| 87 | E | 0.097 |
| 90 | K | 0.075 |
| 91 | Y | 0.281 |
| 93 | R | 0.093 |
| 94 | Q | 0.064 |
| 97 | A | 0.095 |
| 98 | D | 0.159 |
| 100 | V | 0.112 |
| 101 | N | 0.045 |
| 104 | K | 0.068 |
| 105 | L | 0.099 |
| 119 | L | 0.155 |
| 121 | R | 0.112 |
| 123 | F | 0.032 |
| 138 | N | 0.117 |
| 140 | F | 0.081 |
| 171 | W | 0.071 |
| 176 | E | 0.143 |
| 180 | R | 0.122 |
| 182 | A | 0.035 |
| 187 | T | 0.050 |
| 191 | W | 0.043 |

Table S5: Average change in NetMHCpan-4.1 EL score for experimentally binding peptides under residue substitution for peptides binding to HLA-C*04:03

| Pos | AA | Change |
|---|---|---|
| 8 | L | 0.045 |
| 9 | E | 0.190 |
| 11 | S | 0.216 |
| 13 | G | 0.103 |
| 26 | L | 0.099 |
| 28 | E | 0.143 |
| 30 | H | 0.173 |
| 47 | F | 0.037 |
| 57 | V | 0.135 |
| 70 | D | 0.042 |
| 71 | R | 0.161 |
| 74 | A | 0.065 |
| 77 | T | 0.066 |
| 78 | Y | 0.088 |
| 85 | A | 0.083 |
| 86 | V | 0.110 |
| 89 | F | 0.011 |
| 90 | T | 0.082 |
| 89 | T | 0.082 |

Table S6: Average change in NetMHCIIpan-4.0 EL score for experimentally binding peptides under residue substitution for peptides binding to HLA-DRB1*12:02

#### 1.2 Figures

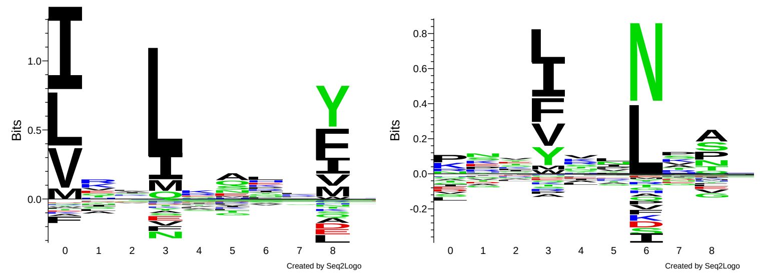


Figure S1: Sequence logos for HLA-DR cores after Gibbs clustering. The left group corresponds to HLA-DRB1*12:02, the right to HLA-DRB3*02:02.

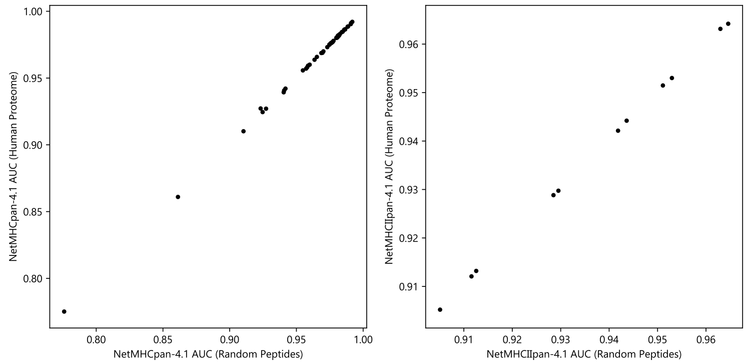


Figure S2: AUC for all alleles when negative control peptides are generated by sampling from the human proteome verses being randomly generated *de novo*.

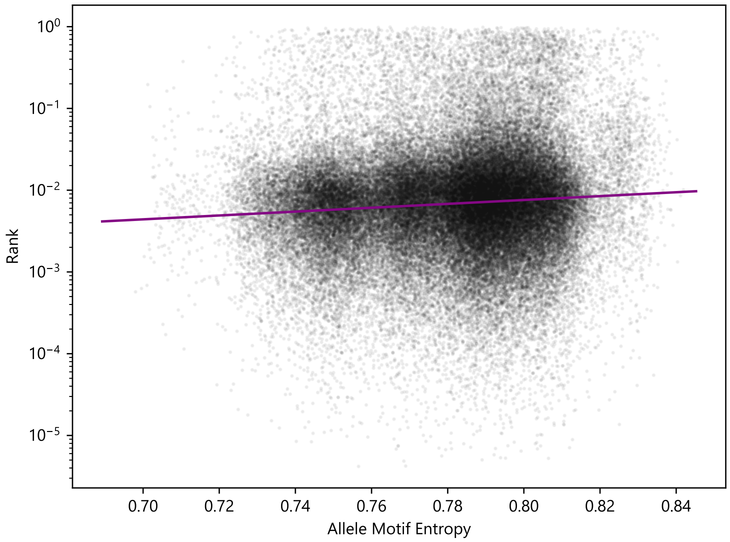


Figure S3: NetMHCpan-4.1 rank predictions vs. allele motif entropy. Rank of the peptides is plotted against the motif entropy of the associated allele. Jittering is performed on the x-axis to allow points to be more easily distinguished.

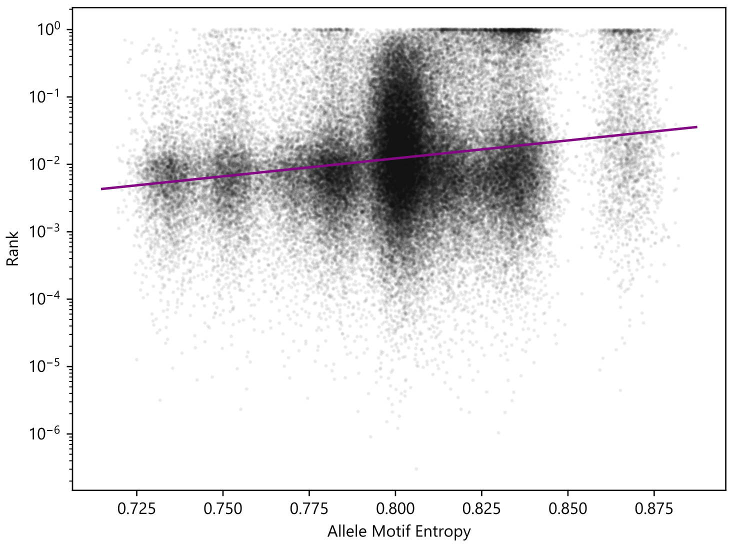


Figure S4: NetMHCIIpan-4.0 rank predictions vs. allele motif entropy. Rank of the peptides is plotted against the motif entropy of the associated allele. Jittering is performed on the x-axis to allow points to be more easily distinguished.

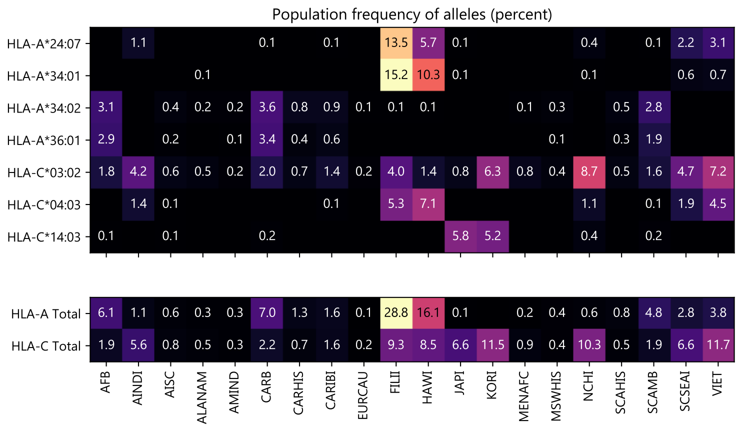


Figure S5: NMDP population frequencies of the alleles from Sarkizova et. al. that are not present in the NetMHCpan training dataset, per 100. Values of less than 0.1% are omitted.

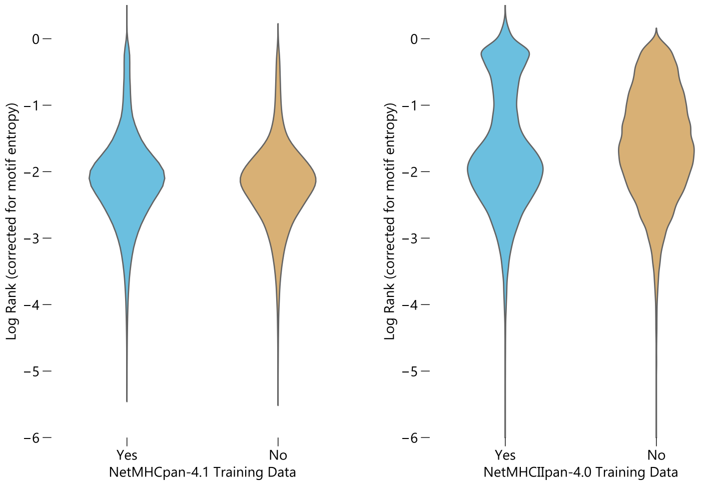


Figure S6: Distribution of entropy-corrected log ranks for peptides corresponding to HLA alleles with and without data in NetMHCpan's training set.

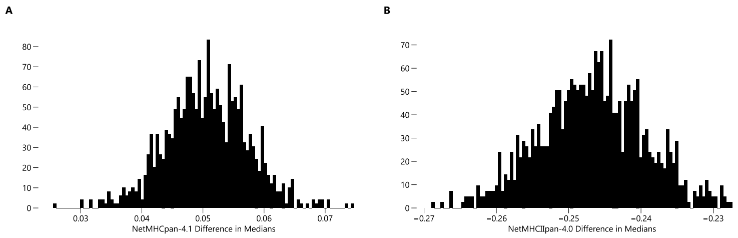


Figure S7: Bootstrap distributions of the median of log ranks of predictions made with training data minus the median of log ranks of predictions made without training data. Positive values indicate the model performs better without training data. (A) Distribution for NetMHCpan-4.1. (B) Distribution for NetMHCIIpan-4.0.

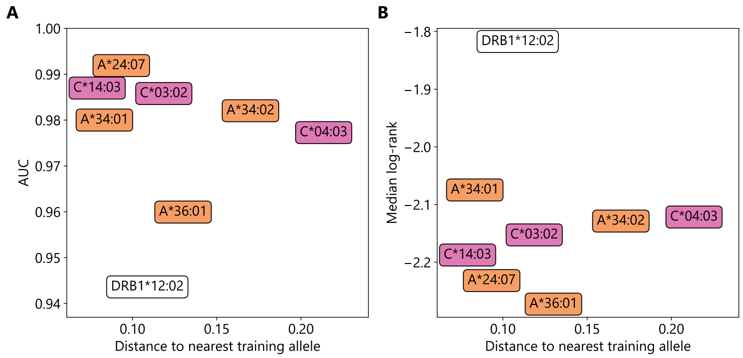


Figure S8: Sequence distance to the nearest MHC allele in NetMHCpan training dataset versus (A) AUC and (B) median log rank of experimentally verified peptides for $n = 8$ HLA alleles without data in NetMHCpan training set. Higher AUC is better, lower median log rank is better. Color corresponds to HLA type.
